# The rhizosphere of *Pappostipa frigida* as a hotspot of active bacterial communities in the Andean steppe of the Atacama Desert

**DOI:** 10.1101/2025.10.03.680286

**Authors:** Daniel E. Palma, Constanza Schapheer, Alexis Gaete, Pamela Aravena, Camila Albarrán-Cuitiño, Constanza Aguado-Norese, Mauricio González, Verónica Cambiazo

## Abstract

**Background:** The rhizosphere is a resource-rich microenvironment where plants, soil nutrients, and microorganisms interact. In arid and semi-arid regions, this tripartite relationship must withstand long periods of drought followed by brief periods of rainfall. In the present study, we employed a combination of RNA/DNA metabarcoding and shotgun metagenomic sequencing to provide insights into the functional capabilities and activity levels of the bacterial communities present in the bulk soil and rhizosphere samples of *Pappostipa frigida*, a grass species endemic to the Andean steppe of the Atacama Desert.

**Results:** The active bacterial community in the rhizosphere of *P. frigida* exhibited greater diversity and a higher Shannon index than the total bacterial community. In terms of beta diversity, the structures of the total and active communities differed markedly between the BS and the RZ. Furthermore, active bacteria in the RZ showed a stronger correlation with total bacterial populations than those in the BS. This finding is consistent with the low proportion of ASVs derived from RNA extractions detected in the BS. Notably, 64% of these putative inactive bacterial populations were identified as active RZ members and 73% grew in culture media, suggesting they were likely dormant. The bacterial communities of the BS exhibited higher abundances of sporulation genes. In contrast, active bacterial communities in the RZ consistently contained higher abundances of genes associated with halotolerance, siderophore synthesis, and resuscitation-promoting factors.

**Conclusions:** The results emphasize the importance of the conditions created by plants in recruiting bacterial populations from the soil and provide insights into how the rhizosphere of arid native plants influences the activity and functional traits of soil microorganisms in their natural habitat. Additionally, this study advances our understanding of the mechanisms employed by soil microorganisms to cope with desiccation in natural environments, establishing *P. frigida* as a model in plant science for studying grass traits and responses to extreme environments.

## Background

Arid regions cover about 45% of Earth’s land surface [1] and are highly sensitive to climate change [2]. In these environments, soil microbiota play a fundamental role in determining the soil properties [3], including nutrient cycling, soil fertility and plant productivity [4, 5], yet their physiological responses to the fluctuation of drought periods under different ecosystems are not well understood [6–8]. In soil bacteria, it has been reported the ability to reduce their internal solute potential, by accumulating sugars, polyols, and amino acids, in order to maintain osmotic equilibrium [9, 10]. In addition, bacteria also could enter a reversible state of reduced metabolic activity, or dormancy when conditions become unfavorable [11, 12]. After the drought period, the rainy season increases the amount of soil water available to microorganisms and causes environmental changes, such as enhancing the availability of soluble nutrient substrates [13]. These changes could induce the transition of bacteria from dormancy to resuscitation through mechanisms involving nutrient-gated ion channels (e.g. GerA) [14] and peptidoglycan hydrolases, such as the resuscitation-promoting factor (Rpf) family [15]. Overall, the entry into dormancy and the transition back to initiate a new cycle of proliferation represent strategies that are particularly important to survive in arid regions [16, 17], where bacterial communities can exist in a large expanse of bulk soil or in association with patches of perennial plants [11, 18, 19]. Since root exudates and plant litter provide abundant and readily available organic matter and micronutrients, several studies revealed that perennial plants growing in arid regions are considered as fertility islands [16, 20]. Therefore, it is possible that the root system can offer a more favorable environment to promote bacterial resuscitation in comparison with their surrounding bulk soil.

The steppe of the Andean Atacama region is characterized by an arid environment (Aridity De Martonne Index = 6.0 [21]) with a short rainy season concentrated in summer (January to March), which induces annual flowering, increases plant richness and biomass, as well as increases soil microbial activity [22]. For the rest of the year (approximately nine months), steppe plants need to cope with prolonged droughts and temperatures ranging from chilling to freezing at night [23].

In this study, we used *Pappostipa frigida*, a dominant, native steppe grass, that shows significant differences in the total bacterial composition between rhizosphere and their surrounding bulk soil [23, 24]. Recently, it has been reported that nutrient availability contributes to the differentiation between bulk soil and rhizosphere bacterial communities of *P. frigida* [25]. Additionally, the rhizosphere of *P. frigida* has been found to contain a variety of primary and secondary metabolites that reliably indicate the environmental conditions of *P. frigida* populations over several years [26] and across an altitudinal gradient of precipitation (10-160 mm yr^-1^) [21]. These metabolites may significantly impact the composition and structure of the rhizosphere microbiome.

In this work, we provide information regarding the functional capabilities of the bacterial communities and their activity levels in the *P. frigida* rhizosphere (RZ) and corresponding bulk soil (BS). We focused on the period immediately after the rainy season. Under these conditions, we hypothesize that (i) there will be differences between the RZ and BS in the abundance of genes associated with survival in these environmental conditions, such as those related to dormancy, resuscitation, and salt tolerance, and (ii) the RZ community of *Pappostipa frigida* will be considerably more active than the BS community due to the relatively more favorable conditions provided by the plant, with this difference being reflected in community structure.

## Methods

### Sampling site and sampling procedure

The sampling site was located at 4,000 m above sea level in the Andes of the Atacama Desert, close to the border between Chile and Argentina (22°55’01.8’’S 67°53’58.0’’W) (Figure 1A). In this place, the soils corresponds to the Entisol soil order [27] and they are dominated by Andean grasslands composed mainly of perennial grasses, including *P. frigida*, one of the plants that had the highest coverage, persistence, and elevation distribution range (3,600 to 4,500 m above sea level) [23, 28, 29] (Figures 1A and 1B). Soil sampling was conducted in April 2022, after the rainy period (January and March). We demarcated a 10 m^2^ area in which samples were taken from five randomly selected *P. frigida* plants of similar sizes and stage of development. Following the protocol described by Fernández-Gómez et al. (2019) [24], we defined two compartments according to their proximity to the roots: the rhizosphere (soil loosely attached to the roots; hereinafter RZ) and the bulk soil (plant-free soil, hereinafter BS). For each plant, the root system was gently shaken to obtain the RZ fraction (25 g) [24, 25]. In addition, BS samples (50 g) were collected at 10 cm depth from the ground and at least 1 m away from each sampled plant. All soil samples (n = 10) were immediately frozen in liquid nitrogen and then kept on dry ice until their arrival at the laboratory, where they were stored at −80°C. Another ten 10 g samples of BS and RZ were stored at 4°C for absolute DNA quantification and for determining soil pH, conductivity, macro-(K, P and Ca) and micro-nutrients (Cu, Fe, Mn and Zn). Additionally, 300 g of a pool of collected BS samples were stored at 4°C to prepare Soil Extract Media (SEM).

**Figure 1.**
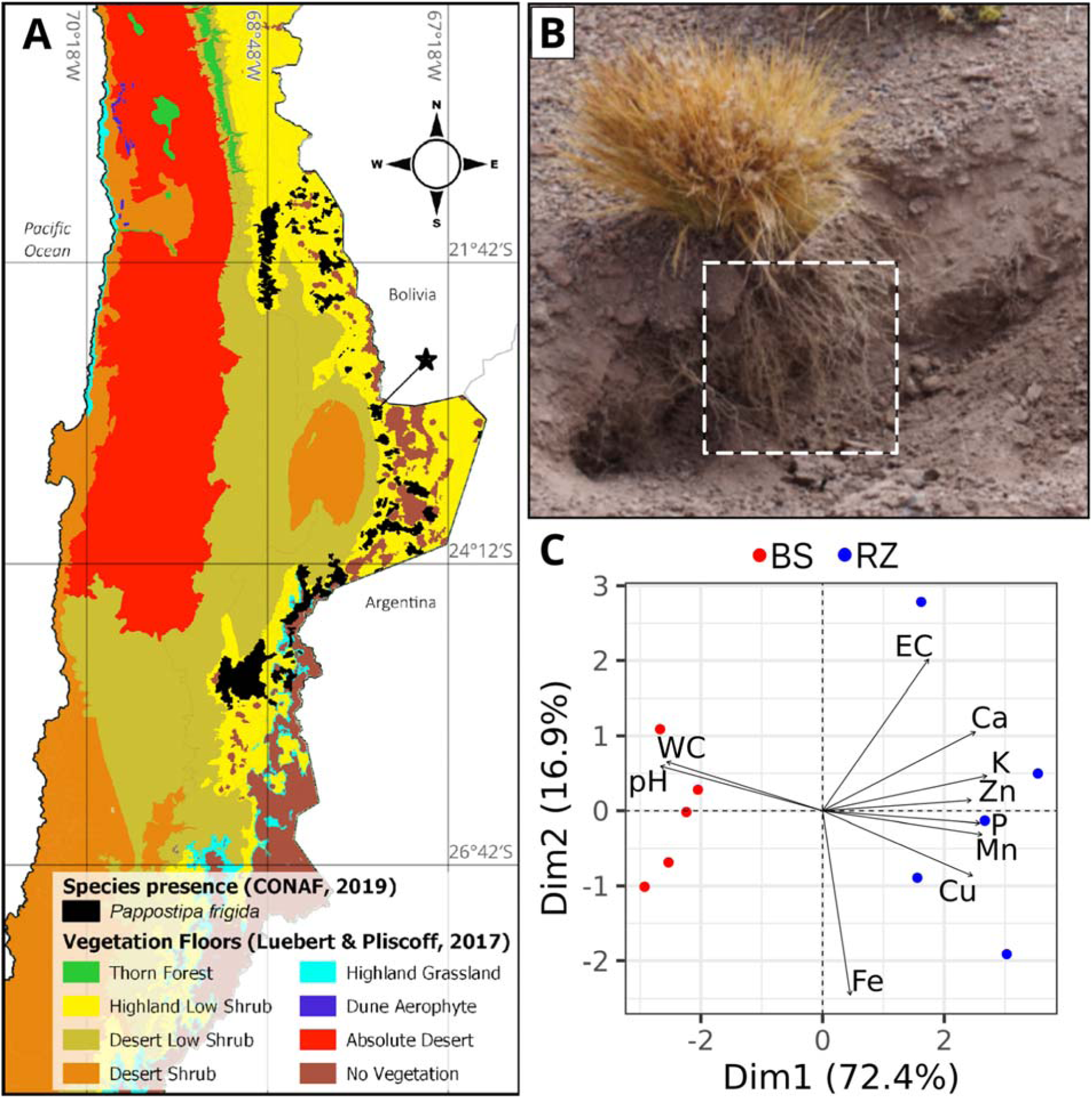
Sampling site and soil properties. (A) Vegetation floors in the northern part of Chile and the distribution of *Pappostipa frigida*. The red star indicates the designated sampling area for *P. frigida*. (B) Photograph of *P. frigida* in the field, with a view of the root system of the plant. The square illustrates the root system of a sampled plant. (C) Principal component analysis (PCA) of soil properties. The points represent the soil samples and the arrows indicate the direction and contribution to the principal components of each chemical property. BS: bulk soil, RZ: rhizosphere, WC: gravimetric water content, EC: electrical conductivity.

### Soil properties

To determine the soluble fraction of metal composition of the soil, the BS and RZ samples were processed as was described by Fernández-Gómez (2019) [24]. Briefly, 1 g of soil was resuspended in 1 mL of distilled water and homogenized for 2 h at room temperature. After mixing, the samples were centrifuged at 11,440 g for 10 min in a Hettich Universal 32R. The soluble fraction was recovered and metals were measured by total reflection X-Ray fluorescence (TXRF) in a Bruker S2 PICOFOX following a previously published protocol [30] and adding gallium as an internal standard element. For soil pH measurement, 2 g of soil were mixed with 5 mL milliQ water and homogenized during 2 h in a S1000 Gyrotwister at 60 rpm. Afterwards, the soil was decanted during 2 h. The pH was measured in triplicate for each sample in a Thermo Scientific Orion 3 star benchtop pH meter [24]. To determine the conductivity of the soil, five grams of soil were diluted in 25 mL of double-distilled water and incubated for 2.5 hours at room temperature under continuous rotation. The samples were then centrifuged at 6000 x g for 10 min, and the supernatant was recovered. Ten mL of the supernatant was used to measure conductivity using a Hanna bench conductivity meter HI 4321. All samples were processed in duplicate. The gravimetric water content of the soil was determined by dividing the water weight (soil fresh weight – soil dry weight) by the soil dry weight after 48 h in an oven at 60 °C when constant weight was reached.

Differences between the soil composition and pH were evaluated using the Wilcoxon rank-sum test corrected for multiple comparisons with the Benjamini-Hochberg method [31]. In addition, a principal component analysis (PCA) was performed using R version 4.2.3 on the centered and scaled physicochemical properties to visualize their influence on the soil samples.

### Soil RNA and DNA isolation

To simultaneously isolate RNA and DNA from soil samples, the PowerSoil total RNA kit (Qiagen) and PowerSoil DNA elution kit (Qiagen) was used following the manufacturer’s recommendations with minor modifications. Briefly, 3 g of soil were used as an initial sample, and the mechanical lysis was performed with a mixture of glass beads and bead solution that was processed for three pulses of 30 s at 6 m/s in a FastPrep-24 (MP Biomedicals). Nucleic acids were quantified by fluorometry using the Qubit dsDNA BR Assay Kit (Invitrogen) for DNA samples and Qubit RNA BR Assay Kit for RNA samples. Integrity of the DNA samples was corroborated in a 1 % agarose gel, while an agarose/formaldehyde gel was used for the RNA samples. The DNA and RNA samples were stored at −20°C and −80°C, respectively.

### Digital PCR analysis of soil bacterial communities

Absolute DNA quantification was performed, in the 10 soil samples, using the QuantStudio 3D Digital PCR system (Thermo Fisher Scientifics, USA), following the manufacturer’s protocols. A commercial TaqMan® probe targeting the 16S rRNA gene was used. For quantification, 2 µL of Absolute Q™ DNA Digital PCR Master Mix (5X), 0.5 µL of the TaqMan® probe-labelled target primer, 1 µL of DNA and 5.5 µL of nuclease-free water were mixed. Subsequently, 9 µL of the reaction mixture was loaded into the QuantStudio™ 3D Digital PCR MAP with the addition of 15 µL of elution buffer. The results were analyzed using the QuantStudio 3D Analysis Suite software. Concentrations of target DNA sequences, together with their Poisson-based 95 % confidence intervals, were calculated using QuantStudio 3D Analysis Suite Cloud.

### Culture of soil bacteria

A Soil Extract Media (SEM, a minimal nutrient medium) [32] and the rich medium Luria-Bertani (LB), which was supplemented with NaCl (LB 10 % NaCl), were used to obtain the culturable microbiota from the BS and RZ samples. SEM was prepared with a combination of 300 g of soil and 300 mL of distilled water homogenized by stirring for 4h. Subsequently, the mixture was centrifuged at 9,000 rpm for 10 min, the supernatant was recovered, and 1.5 % agar was added, autoclaved, and plated on petri dishes. LB 10 % NaCl (10 g of tryptone, 100 g of NaCl, 5 g of yeast extract, 15 g of agar and 1 L of distilled water) was sterilized by autoclaving and plated on petri dishes. For culture assays, a suspension of 1 g of each soil sample in 1 mL of 1X Phosphate-Buffered Saline (PBS 1X) was used as inoculum. The mixture was homogenized by shaking in a Revolver Rotator (LabNet) for 2 h and then decanted by gravity for 30 min at room temperature. After that, 100 µL of supernatant were inoculated into SEM and LB 10 % NaCl plates and distributed with an L-shaped cell spreader. A total of 10 plates (five BS samples using SEM, and five BS samples using LB) were incubated at 30°C for 72 h, and after this time, the plates were scraped to recover all bacterial growth. Bacteria were suspended in 200 µL of sterile PBS, homogenized by vortexing and used for DNA isolation using the DNeasy Blood & Tissue kit (Qiagen), following manufacturer’s recommendations. The 10 DNA samples obtained were quantified by fluorometry using Qubit dsDNA BR Assay Kits (Invitrogen), their integrity was corroborated in a 1 % agarose gel and used for total 16S rRNA metabarcoding.

### 16S rRNA metabarcoding

DNA and cDNA samples were prepared for sequencing by Molecular Research LP (Mr. DNA Lab). Briefly, the RNA was pretreated with DNase I (RNase-free) (New England Biolabs) and reverse transcribed using the High-Capacity RNA-to-cDNA kit (Applied Biosystems). DNA and cDNA samples were amplified using primers 28F (5’-GAGTTTGATCNTGGCTCAG-3’) and 519R (5’-GTNTTACNGCGGCKGCTG-3’), targeting the V1-V3 region of the bacterial 16S rRNA gene, and the HotStarTaq Plus Master Mix Kit (Qiagen) for sequencing on an Illumina MiSeq platform. The reads were processed using Qiime 2 v2022.2 plugins [33]. The barcodes and primers were removed from the demultiplexed reads using the Cutadapt [34] *trim-paired* plugin. ASVs (Amplicon Sequence Variants) were generated using the DADA2 [35] *denoise-single* plugin and were assigned a taxonomy using the *classify-sklearn* plugin with a naive Bayes classifier trained on the V1-V3 region of the 16S rRNA sequences from the GTDB r226 database [36] downloaded using the RESCRIPt plugin [37]. Only ASVs that met any of the next conditions were retained: i) detected in the soil samples by DNA and RNA sequencing, or ii) detected in soil and culture samples. Additionally, to identify spore-forming ASVs, the KEGG Orthology gene groups (KOs) [38] associated with each ASV were predicted using PICRUSt2 [39]. An ASV was designated as spore-forming if it belonged to the Bacillota phylum and possessed a minimum of 20 of the KOs associated with sporulation genes (see Table S1 for details).

For diversity analyses, bacterial ASVs samples were rarefied to 58,000 counts. The R package vegan v2.6-4 [40] was used to calculate the alpha diversity indices (richness and Shannon index). The statistical significance of the comparisons of alpha diversity indices between groups was evaluated using the Wilcoxon rank-sum test. For beta diversity analysis, a phylogenetic tree was constructed using FastTree v2.1.11 [41] with the gtr model based on an ASV alignment generated with Infernal v1.1.4 [42] (using the cmsearch command with gathering cutoffs) and the Rfam [43] bacterial small subunit ribosomal RNA covariance model (RF00177). Weighted UniFrac distances were then calculated from this tree using the R package phyloseq v1.42 [44].

### Shotgun metagenomic sequencing

Library preparation and shotgun metagenomic sequencing were performed using the Illumina NovaSeq 2 × 250 bp platform at Molecular Research LP (Mr. DNA Lab), following the manufacturer’s guidelines. Illumina reads were trimmed using fastp v0.23 (--cut_front --cut_right --length_required 30) [45].

Additionally, for both soil types (BS and RZ), DNA from the five corresponding samples was pooled and sequenced using Oxford Nanopore Technologies (ONT). Library preparation was performed using genomic DNA that had been previously fragmented to an N50 of 8,000 bp with a g-TUBE (Covaris) and subsequently processed with the Ligation Sequencing Kit SQK-LSK109 (Oxford Nanopore Technologies) according to the manufacturer’s protocol. Sequencing was carried out on a MinION device (version 1B) (Oxford Nanopore Technologies) with an R9.4.1 flow cell. Basecalling was performed using Guppy v6.5.7 (Oxford Nanopore Technologies). Reads were retained only if they had a minimum mean Q-score of 7 and a minimum read length of 1,000 bp.

For each soil type (BS and RZ), Illumina and Nanopore reads were co-assembled using metaSPAdes v4.1.0 in hybrid mode [46, 47]. Illumina reads were mapped to the contigs using BWA-MEM v0.7 [48], and the abundance of each contig was calculated with CoverM [49] using the mean coverage method. Coding sequences (CDSs) were predicted using Pyrodigal v2.1 [50], a Python library binding to Prodigal [51]. Functional annotation of the proteins, including KEGG KO assignment [38], was performed using eggNOG-mapper v2.1.9 [52]. Finally, the abundance of each selected KEGG KO (Table S1) was calculated as the sum of the abundances of the contigs containing the corresponding CDSs, and a differential abundance analysis was carried out using LinDA [53] from the R package MicrobiomeStat v1.2 in proportion mode.

## Results

### Abiotic properties of the RZ and the BS

A general description of the BS and RZ abiotic properties was performed through analysis of their physicochemical characteristics and the composition of their soluble metal fractions (Table S2). The comparative analysis demonstrated that, with the exception of soluble iron, which did not exhibit a significant difference between the compartments, there was a substantial increase in electrical conductivity and in the elements K, P, Ca, Cu, Mn, and Zn in the RZ compared to BS (Wilcoxon rank-sum test; p ≤ 0.05). In contrast, a significant decrease in pH and water content was observed in the RZ compartment (Wilcoxon rank-sum test; p ≤ 0.05). The principal component analysis (Figure 1C) indicated that these variables were highly correlated with the first principal component, which explained 72.4% of the variance and separated the BS samples from the RZ samples.

### Functional analysis of soil metagenomes

The total number of Illumina reads obtained for BS and RZ samples was 3.1 ± 0.7 and 3.5 ± 0.4 Gbp per sample, respectively, from shotgun sequencing. For Nanopore sequencing, the total number of reads obtained was 9.0 and 7.3 Gbp for BS and RZ, respectively. The metagenome assembly of BS by employing reads from both strategies yielded 5.5 × 10^6^ contigs and 2.8 Gb of total length, with an N50 length of 1,086 bp. In the case of RZ, the metagenome consisted of 6.9 × 10^6^ contigs and a total length of 3.3 Gb (N50 length of 902 bp). A total of 4.9 × 10^6^ and 5.8 × 10^6^ prokaryotic proteins were annotated within BS and RZ assembled contigs, respectively.

In the BS and RZ metagenomes, we explored the patterns exhibited by traits reported as relevant to the survival of microorganisms under arid and semi-arid conditions (Figure 2, Table S3). In order to do that, we measured the relative abundance of genes associated with dormancy mechanisms, salt tolerance and iron cycling based on siderophore synthesis in BS and RZ microbial communities.

**Figure 2.**
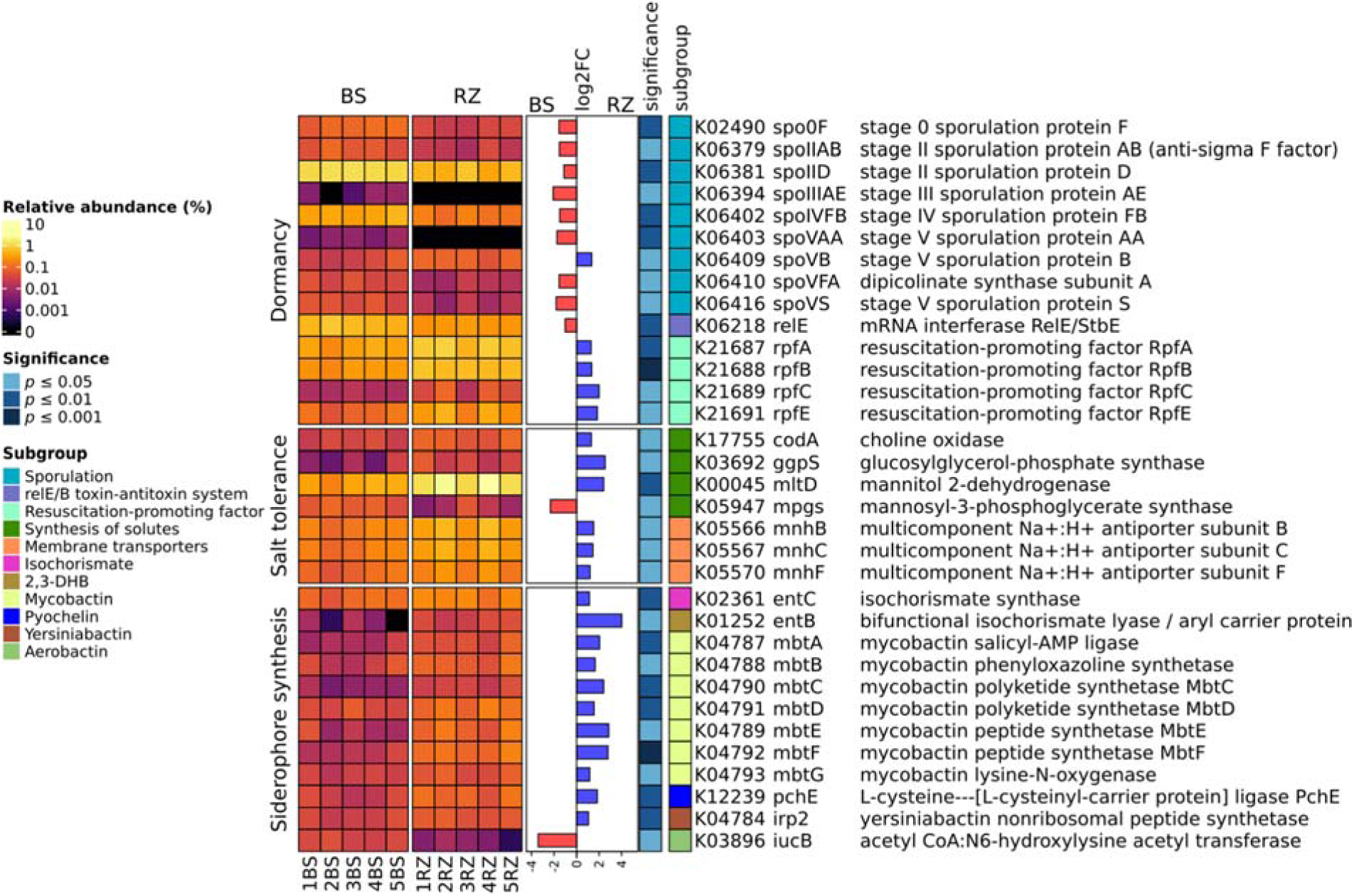
Functional potential of the soil samples. Differences in abundance of the KEGG KOs between bulk soil (BS) and rhizosphere (RZ). Only KEGG KOs with an absolute log_2_ fold change (FC) greater than 1 and a *p*-value less than 0.05 are displayed.

In particular, genes implicated in sporulation exhibited a significantly higher abundance in BS (Figure 2; |log_2_FC| ≥ 1, *p ≤* 0.05). With respect to dormancy-related toxin-antitoxin genes, only *rel*E showed increased relative abundance in BS. In contrast, four genes encoding dormancy resuscitation factors (*rpf*A, *rpf*B, *rpf*C and *rpf*E) were found to be significantly more abundant in RZ (Figure 2; |log_2_FC| ≥ 1, *p ≤* 0.05), suggesting that bacteria inhabiting the *P. frigida* rhizosphere have different strategies for entering dormancy and transit to resuscitation compared to bacteria living in bulk soils. Furthermore, a significant increase in the abundance of genes implicated in salinity tolerance was observed in RZ, including Na^+^ efflux-related genes (*mnh*B, *mnh*C, *mnh*F; |log_2_FC| ≥ 1, *p ≤* 0.05) and organic solute biosynthesis (*cod*A, *ggp*S, *mlt*D; |log_2_FC| ≥ 1, *p ≤* 0.05).

Considering that siderophores facilitate the resuscitation of stressed and viable but non-cultivable cells [54, 55], and also increase soil iron availability under saline conditions [56], the relative abundance of genes implicated in the synthesis of siderophores was also ascertained. The results indicated that 11 genes encoding for enzymes involved in the synthesis of isochorismate (*entC*), 2,3-dihydroxybenzoate (*entB*), mycobactin (*mbt* genes), pyochelin (*pchE*) and yersiniabactin (*irp2*) showed a significant increase of their relative abundance in RZ (Figure 2; |log_2_FC| ≥ 1, *p ≤* 0.05). The results demonstrated that, at the community level, the majority of the tested genes associated with resuscitation mechanisms, salt tolerance, and siderophore production that exhibited significant differences were more abundant in the RZ compartment. In contrast, the BS compartment exhibited a greater representation of genes associated with sporulation.

### Analysis of total and active communities

Given the significant differences in traits associated with the survival of microorganisms between BS and RZ, our next step was to assess whether these were accompanied by changes in the composition of the total and active members of the soil bacterial communities. To that end, we generated amplicon sequence variants (ASVs) from the amplification of 16S rRNA sequences derived from DNA and RNA extractions. The primary benefit of employing ASVs in this scenario is that they facilitate the efficient identification of shared individuals across diverse samples. Following the definition given by [12], the ASVs of 16S rRNA genes from DNA extractions were considered to be a representation of the total bacterial community, including active, dormant and dead individuals [57]. Meanwhile, the ASVs of 16S rRNAs from RNA extractions were considered to be representative of active members of the soil bacterial community.

A total of 4,626 and 3,783 ASVs (Figure 3A, Table S4) were identified in the DNA and RNA samples, which were classified into 810 and 718 genera, respectively (Table S5). The percentage of ASVs detected exclusively in the RZ was higher in RNA samples (60 %) than in DNA samples (31 %) (Figure 3A). In contrast, in both RNA and DNA samples only a small percentage of ASVs (11 % and 1 %) were exclusive to the BS compartment (Figure 3A).

**Figure 3.**
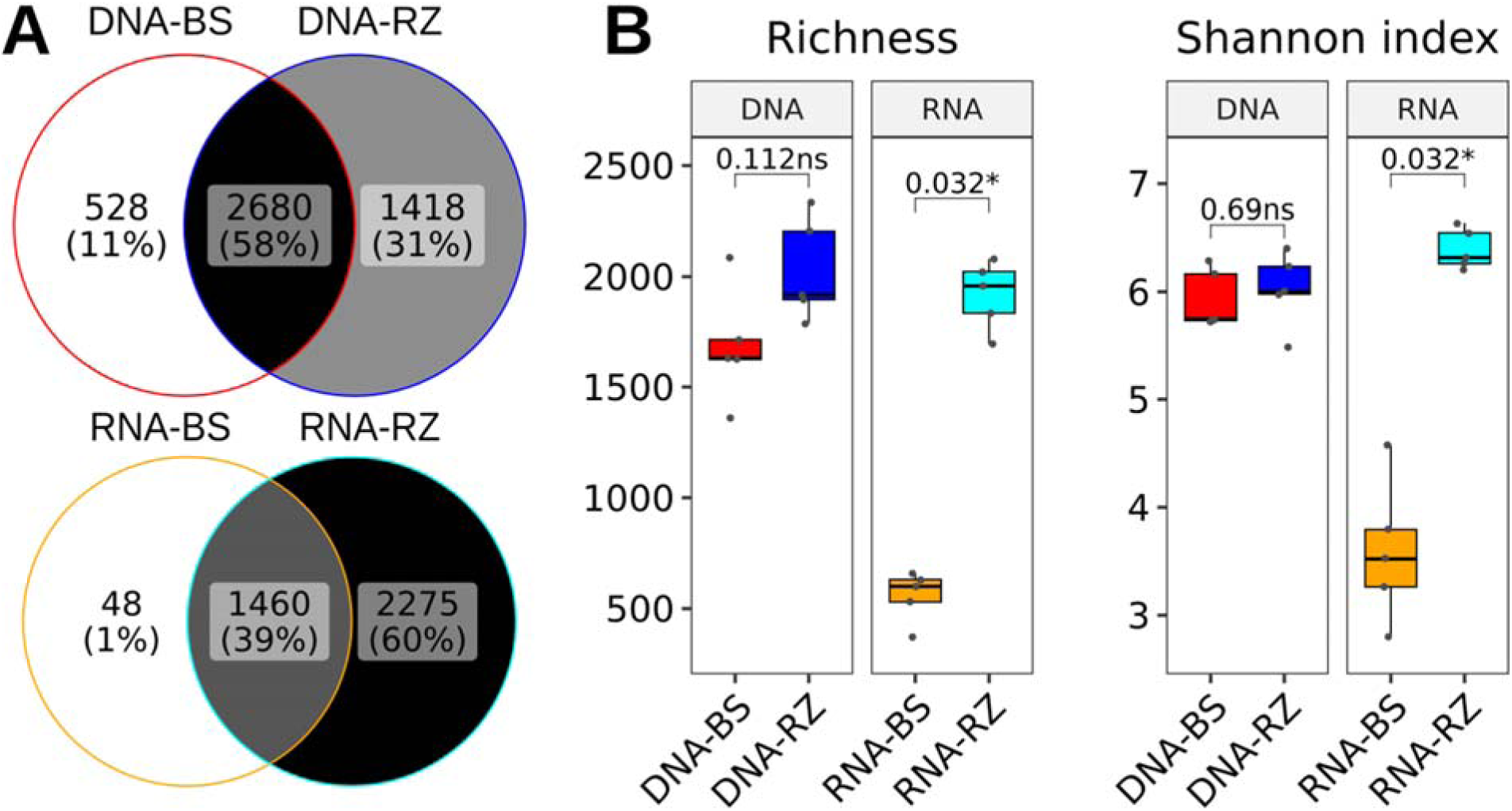
Alpha diversity metrics of the total (DNA) and active (RNA) bacteria present in the soil samples. (A) Venn diagram illustrating the detection of ASVs in bulk soil (BS) and the rhizosphere (RZ). (B) Boxplots depicting the alpha diversity metrics (richness, Shannon index) at the ASV level. ns: not significant, * p < 0.05.

Rarefaction curves indicated that at a depth of 58,000 counts, the alpha diversity metrics neared a plateau, with only minimal increases observed beyond this point (Figure S1). Using this cut-off, we observed a considerable difference in alpha diversity metrics between total and active communities when comparing BS and RZ (Figure 3B). While the total communities (DNA) in BS and RZ showed no significant differences in richness or Shannon index, with median richness values of 1,632 and 1,918, and median Shannon index values of 5.75 and 6.00, respectively, the active community (RNA) in BS had significantly lower richness and Shannon index than in RZ (*p ≤* 0.05), with median richness values of 600 and 1,958, and median Shannon index values of 3.52 and 6.32, respectively.

To evaluate the differences in beta diversity, we calculated the weighted UniFrac distances between the samples, tested for group differences using PERMANOVA, and generated a complete-linkage hierarchical clustering. The results showed that the 20 samples (10 from DNA and 10 from RNA) clustered into two main groups (Figure 4A) that separated the bacterial communities of RZ from BS, revealing a greater similarity between bacterial communities from the same soil compartment, regardless of whether they were derived from RNA or DNA. It is interesting to note that, while there was no significant difference in the total and active communities in the RZ group, DNA-derived samples were tightly clustered and significantly different (PERMANOVA, *p ≤* 0.05) from RNA-derived samples in the BS group. (Figure 4A).

**Figure 4.**
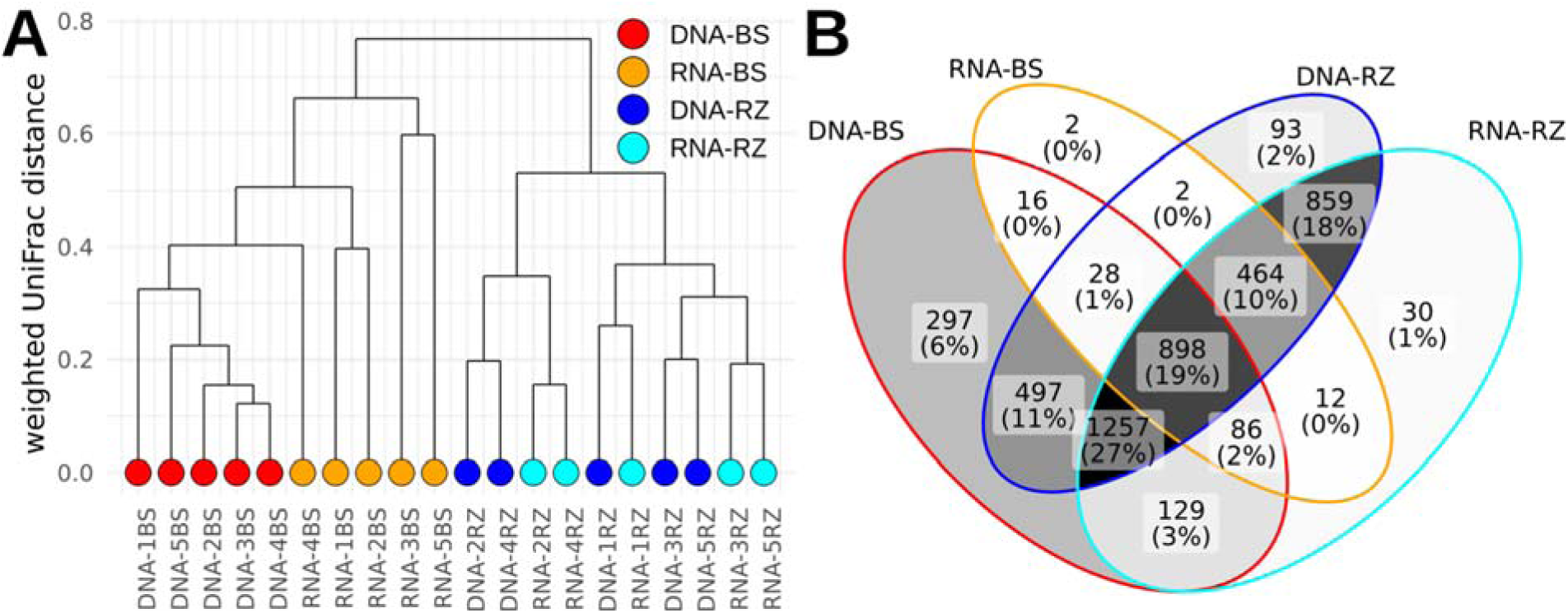
Beta diversity analysis of the total (DNA) and active (RNA) bacteria present in the soil samples. (A) Complete-linkage hierarchical clustering constructed from weighted UniFrac distances between samples. (B) Venn diagram illustrating the ASVs detectable in each sample type. BS: bulk soil, RZ: rhizosphere.

Of the 4,672 ASVs and 818 genera detected in any of the soil sample types, 898 ASVs (19 %) (Figure 4B, Table S4) and 317 genera (39 %) (Table S5) were shared across all sample types. On the other hand, considering both active and total ASVs, a total of 982 ASVs (21 %) and 113 genera (14 %) were found exclusively in RZ communities, while 315 ASVs (6.7 %) and 40 genera (4.9 %) were found exclusively in active and total BS communities. Interestingly, 1,386 (64 %) of the 2,180 ASVs detected by DNA but not by RNA in BS were detected in the active RZ community (Figure 4B), suggesting that differences of physical and nutritional features of root system could regulate the taxonomic composition of rhizosphere by mediating microbial activities of shared members among BS and RZ. This fraction contributed with 11.1% of the relative abundance of the active RZ community (Table S4). The observed dissimilarity between active and total BS communities was also evident when comparing the number of genera per phylum, since most phyla had fewer genera detected in the active community (n = 375) than in the total community of BS (n = 657) (Figure S2), including Actinomycetota (110 in DNA-BS, 81 in RNA-BS), Pseudomonadota (139 in DNA-BS, 104 in RNA-BS) and Acidobacteriota (62 in DNA-BS, 42 in RNA-BS) (Table S5). In contrast, in RZ, the number of detected genera for each phylum tends to be more similar between total and active communities. This is evident in Actinomycetota (134 in DNA-RZ, 127 in RNA-RZ), Pseudomonadota (176 in DNA-RZ, 171 in RNA-RZ) and Acidobacteriota (66 in DNA-RZ, 65 in RNA-RZ). When considering both DNA and RNA, it was found that relatively few genera were exclusive to BS (n = 40) or RZ (n = 113), and together they accounted for less than 1% of the mean relative abundance in their respective sample types (see Table S5 for details). Therefore, under these conditions, the majority of the ASVs (Figure 4B) and genera detected in BS were also present in RZ, albeit with differences in their abundances, as suggested by the beta diversity analysis (Figure 4A).

### Relationship between the relative abundances of the active and total bacterial communities in the BS and the RZ

In the total BS community, we were able to detect genera that were not detected in the RNA samples, regardless of their abundance in the DNA-derived samples (Figure 5A, arrow), while in the RZ (Figure 5B) the few genera that were not detected in the RNA samples corresponded mainly to those with the lowest abundance in the DNA-derived community. In addition, a Spearman’s rank correlation test showed that, when comparing the abundance of each genus in the total community with that in the active community, both BS and RZ displayed a significant monotonic positive association (*p ≤* 0.05). However, the RZ had a considerably higher ρ (rho) value (0.74) than the BS (0.28). Furthermore, it was noted that the predominant phyla (Actinomycetota, Pseudomonadota and Acidobacteriota) were present across the full spectrum of abundances in the active and total BS and RZ communities (Figures 5A and 5B). However, the phylum Bacillota (Figure 5, green dots) was an exception, with most of its highly abundant genera not detectable in the active community of the BS or showing a relatively low abundance in the RZ (Figure 5, green dots). The differences observed in the taxonomic composition and proportion of the total and active bacterial populations of the RZ compared to those of the BS were accompanied by a significant increase in total bacterial abundance as measured by digital PCR, which increased from 1,845.6 (± 120.7) copies/ng of DNA in the BS compartment to 2,657.6 (± 96.2) copies/ng of DNA in the RZ compartment (see Table S2).

**Figure 5.**
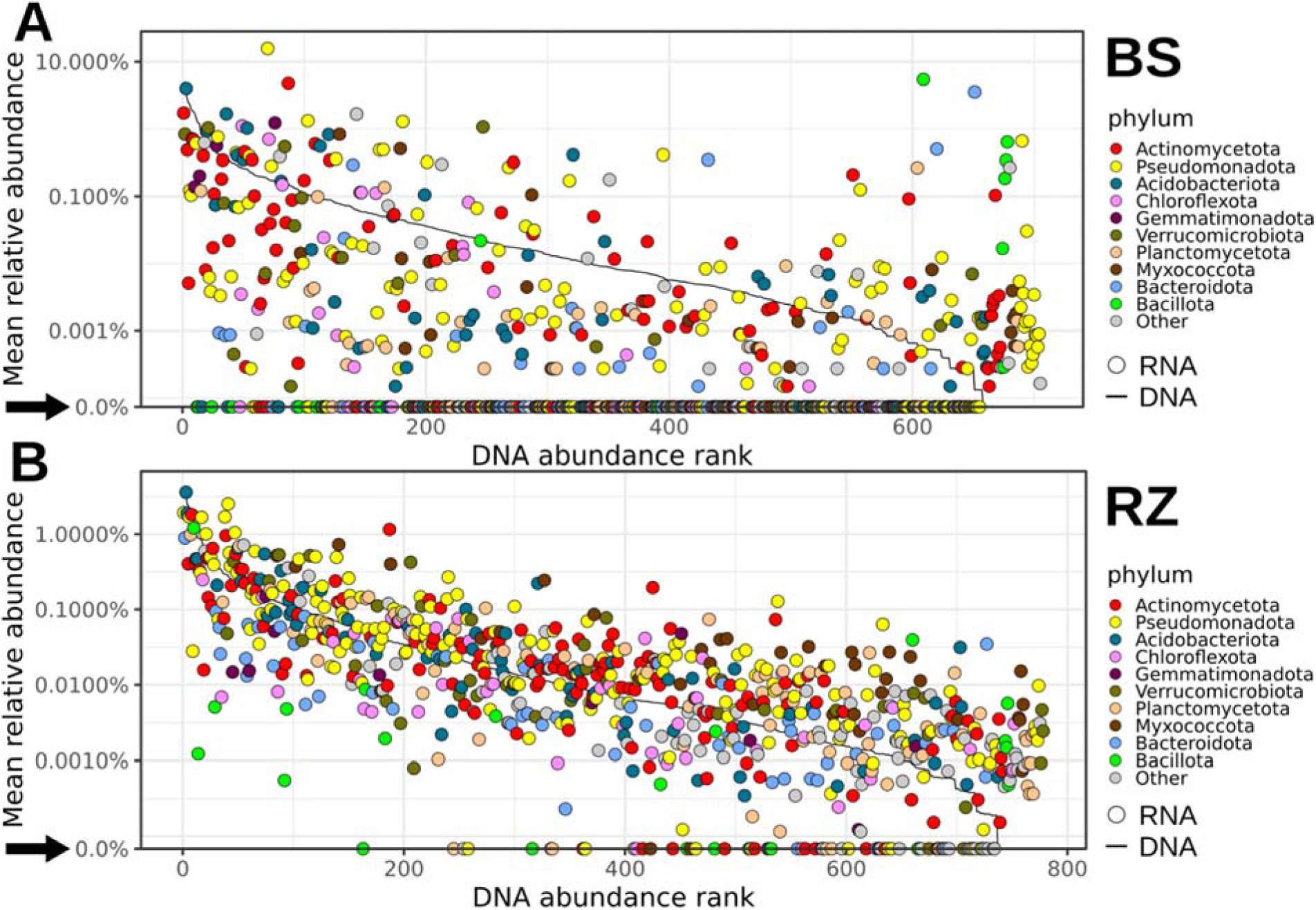
Relationship between the mean relative abundances of DNA and RNA for genera detected in the bulk soil (BS) and the rhizosphere (RZ). Genera are ordered by their DNA abundance rank and are colored according to their phylum. The relative abundance of DNA is represented by a line, while the relative abundance of RNA is shown as dots for the genera present in (A) BS and (B) RZ. The black arrows indicate the genera without detectable RNA abundance.

### Reactivation of potential dormant bacteria from BS and RZ by *in vitro* assays

A culture-based approach, in combination with 16S rRNA gene amplicon sequencing, was employed to investigate whether some of the inactive bacteria at the BS compartment (see Figure 6A, arrow) could be detected in these soil samples cultivated for 72 hours under controlled laboratory conditions. For this purpose, we used Soil Extract Media (SEM), a culture medium made with the soluble elements of the soil, and LB medium supplemented with 10% NaCl, as salt content has been previously reported as a key factor in shaping the Atacama soil microbiome [32, 58–60].

**Figure 6.**
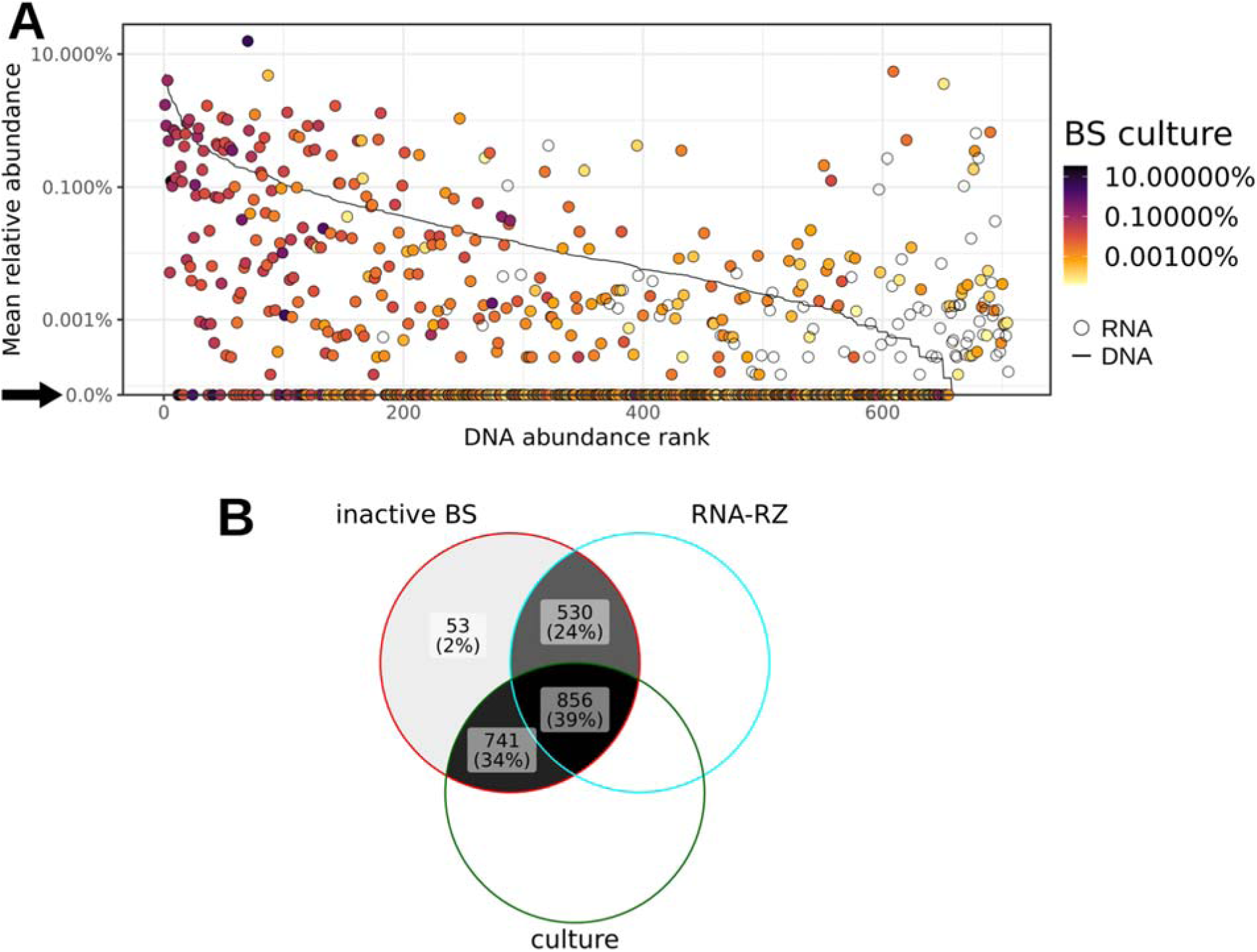
Detection of soil bacteria in cultures. (A) Relationship between the mean relative abundances of DNA and RNA in the soils, and DNA in the cultures, for genera detected in bulk soil (BS). Genera are ordered by their DNA abundance rank and are colored according to their mean abundance in the respective cultures. The relative abundance of DNA in the soil is represented by a line, while the relative abundance of RNA is shown as dots. The black arrow indicates the genera without detectable RNA abundance. (B) Inactive ASVs from BS shared with the RNA-RZ and culture (BS-SEM and BS-LB 10% NaCl) samples.

Our results indicated that of the 2,180 ASVs detected as inactive in BS (Figure 6B), 769 ASVs (35.3%) were recovered in the cultivable fraction grown on SEM and 1,367 ASVs (62.7%) were recovered from LB 10% NaCl cultures (Table S4). Together, these represented more than 70 % (1,597) of the total inactive ASVs in BS (Figure 6B). Upon integrating the data from *in vitro* assays with that of the active RZ bacterial community, we identified 856 ASVs that were inactive in BS and detected in both the active RZ community and the cultures (Figure 6B). Finally, among the inactive ASVs predicted by PICRUSt2 to belong to spore-forming Bacillota, we found that of the 92 ASVs, 17 were detected by RNA in RZ, while 87 were recovered from cultures, and 14 were present in both conditions (Figure S3, Table S4). Among these 92 ASVs, the most abundant genus in the LB 10% NaCl cultures was *Priestia*, representing 67.8% of the average abundance, while in the SEM the most abundant genus was *Paenibacillus_E* with 14.2%. However, neither of them showed substantial abundance in the active fraction of the RZ.

## Discussion

### Functional capacities of the bacterial communities of the BS and the RZ: dormancy, salt tolerance and siderophore synthesis traits

Analysis of shotgun metagenomic data revealed significant differences in microbial abundance and function between BS and RZ compartments. The genes that demonstrated higher consistency in their abundance in RZ were those implicated in the synthesis of siderophores, a process that has the capacity to regulate the bioavailability of iron and to protect against phytopathogens [61]. Some of these genes include *entC* and *entB*, which are involved in the production of isochorismate and 2,3-dihydroxybenzoate, precursors of multiple siderophores [62]. Furthermore, a greater abundance of several *mbt* genes involved in the synthesis of mycobactin was observed in RZ compared to BS. This siderophore is typically produced by *Mycobacterium* species, with some species having been isolated from plants and their rhizosphere [63–65]. Mycobactin was also detected in a strain of *Nocardia mangyaensis* that was isolated from an endolithic environment, suggesting that mycobactin siderophores in *Nocardia* might play a role in enabling survival under extreme conditions [66].

It has been reported that bacteria in the Atacama Desert soil survive by adjusting their physiological metabolism within a given range of salinity fluctuations in order to respond to salt stress [32, 58–60, 67]. Consistently, we found that the genes mnh, encoding components of an Na+/H+ antiporter, which are a common mechanism used by bacteria to maintain intracellular sodium homeostasis, were more abundant in the RZ compartment [68, 69]. As part of the bacterial osmoadaptation strategy, genes involved in synthesis of osmoprotectants were significantly more prevalent in the RZ compartment. One of these genes is *codA*, which encodes choline oxidase. This enzyme is found in certain soil bacteria, such as *Arthrobacter pascens*, and it catalyzes the conversion of choline to glycine betaine [70, 71] an important osmoprotectant in bacteria [72]. Other genes include *ggps*, *mltD*, and *mpgs*, which are involved in the biosynthesis of the organic solutes glucosylglycerol, mannitol, and mannosylglycerate, respectively [73–75]. These compounds accumulate in microbial cells in response to osmotic stress [76].

Additionally, microorganisms can enter dormancy to cope with unfavorable environmental conditions. Thus, it has been estimated that only about 1% of the soil microbial biomass is metabolically active [77]. The remaining viable biomass serves as a reservoir of dormant microorganisms that plays a crucial role in population persistence, biodiversity maintenance, and ecosystem functioning [78–80]. In this work, we found higher abundance of genes associated with sporulation in the BS, which is consistent with meta-analysis showing that strategies based on spore formation were more abundant in bulk dryland soils [81]. The formation of endospores is a mechanism unique to the Bacillota phylum that allows its members to survive adverse conditions [82]. We found that some spore-forming Bacillota genera, such as *Priestia* and *Paenibacillus_E*, were inactive in the BS but grew extensively in the culture media. However, in the active community of the RZ, these genera generally exhibited low activity levels compared with other phyla. In other words, they are capable of responding to favorable conditions, but this trait does not appear to provide them with an advantage in dominating the rhizosphere of an established plant. However, it would be interesting to examine whether they are more capable of responding to changes in resource availability at earlier stages of plant development [83]. In addition, we found that resuscitation-promoting factor genes were more abundant in the RZ compartment. These factors, found in the phyla Actinomycetota and Bacillota, facilitate peptidoglycan hydrolysis, which helps resuscitate dormant bacteria with thick cell walls that withstand adverse conditions [84]. Actinomycetota was one of the most abundant phyla in our soils, and its genera showed activity levels in the RZ consistent with their abundance in the total community.

### Diversity of the total and active bacterial communities of bulk soil and rhizosphere

Given the pivotal role of active bacteria in driving biogeochemical processes within the soil matrix and interacting with the roots of plants [85], it is necessary to assess not solely the alterations in the total community, but also the degree of activity among the community members. In particular, unfavorable conditions, such as prolonged drought in arid regions [86], have been shown to result in an increase in dormant bacteria [12, 79]. In addition, such conditions can also lead to cell death and the generation of relic DNA, which can be detected in metagenomic experiments [57, 87]. In the present study, RNA sequencing was employed as an indicator of bacterial activity, given that RNA is exclusively synthesized by actively growing cells and rapidly degrades [88]. The sampling was conducted following a brief period of rain, a time frame that maximized the probability of identifying active bacteria in both the BS and the RZ. This approach enabled the assessment of whether the nutritional contribution from plant litter and exudates could enhance the diversity of active bacteria at the RZ in comparison to the BS.

Our findings indicate that the active bacterial community from RZ samples exhibited a significantly higher diversity (as measured by richness and Shannon’s index) compared to BS samples. In contrast, total bacterial communities showed no significant differences in alpha diversity indices between RZ and BS. The most probable cause of the observed diversity differences between active members of *P. frigida* from BS and RZ is the distinct soil conditions found in each location. In line with this observation, a higher nutrient concentration was detected in RZ compared to BS. This increase in nutrient availability corresponds with a higher proportion of the RZ bacterial community that is active, which could be associated with a change in composition and with the soil biomass [89–91]. This explains the higher correlation between RNA and DNA abundances in RZ (Spearman’s ρ of 0.74) than in BS (Spearman’s ρ of 0.28) and the significant increase in absolute DNA quantified in the RZ. In this regard, a previous report indicates that the RNA:DNA ratio is significantly lower in Calcisol, a soil with relatively limited nutrient content, compared to Retisol and Luvisol [92]. It is notable that the addition of glucose to these soils resulted in an increase in the RNA:DNA ratio of Calcisol to levels that exceeded those observed in other soil types, which remained unaltered.

Given that the majority of ASVs identified in the DNA samples were shared between BS and RZ, and that the majority of ASVs not detected by RNA in BS were detected by RNA and DNA in RZ, it is highly probable that most of these individuals were in a dormant or deceased state in BS. The present findings indicate that the stimulation of a bacterial community in response to elevated nutrient availability within the RZ compartment may constitute a viable strategy for the cessation of dormancy in soils with comparatively limited nutrient content [77]. Indeed, under the conditions of this study, it was found that of the 2,180 inactive ASVs at the BS, 1,386 were detected as active ASVs at the RZ. This result suggests that the majority of the inactive ASVs in the BS at the Andean steppe could become reactivated in response to nutritional conditions derived from root exudation and serve as a source of inoculum for the rhizosphere bacterial community of *P. frigida*. In this regard, a previous study [25] revealed that a subset of the ASVs exhibited a positive response to enhanced iron availability, increasing their abundance and thereby reducing the disparities between the total bacterial communities present in the BS and the RZ compartments.

Therefore, it can be argued that the results of the present study are consistent with the model in which plant fertility facilitates bacterial survival during drought periods at the RZ [16] and/or the transition of inactive bacteria into active states [93], thereby promoting the establishment of these microbial populations, which would otherwise remain inactive in the oligotrophic environment of the BS [83, 92]. This observation indicates that the active bacteria found associated with the *P. frigida* root system may have originated from the same original bacterial community present in the BS. The data concerning the taxonomic composition of the active fraction of BS and RZ of *P. frigida* suggests that the rhizosphere activates a fraction of the inactive microbial community in BS, which has the capacity to respond to changes in resource availability. This, in turn, can modify the resource conditions of plant root systems. Alternatively, the root system may enable bacterial survival during drought periods due to its more favorable nutritional conditions.

On the other hand, beta diversity analysis revealed that the total and active bacterial populations in BS were separated into different groups. According to the microbial dormancy model proposed by Jones and Lennon [12], this dissimilarity pattern could be interpreted as the result of a combination of high dormancy and strong environmental cues producing differences in structure between total and active communities. The presence of a single cluster mixing total and active bacterial populations in the RZ suggests that, in contrast to the BS, most bacteria remain metabolically active in the RZ, possibly due to the reactivation of dormant individuals recruited from the BS. Considering the adverse conditions of the Andean steppe, these differences between the BS and the RZ could result from differences in soil nutrient availability. To examine the reactivation of dormant bacteria in BS communities further, we used two previously defined culture media [32] that were effective in isolating soil bacteria from the same area [94]. In combination with high-throughput sequencing [95], we observed that 62% (856 ASVs) of the active ASVs in the rhizosphere that were detected as inactive in the bulk soil (1,386 ASVs) could grow in SEM and/or the LB 10% NaCl medium. These results support the possibility that at least some of the inactive fraction in the BS represents a dormant fraction of the bacterial community that could be recruited by the root system of the plant *P. frigida*.

## Conclusions

In this study, we report clear distinctions between bacterial communities in bulk soil and in the rhizosphere of *P. frigida*. These distinctions include taxonomic composition, functional potential, and the distribution of active versus total bacterial communities in these two soil compartments. Our results suggest that the rhizosphere of the Andean steppe grass acts as an oasis for soil bacteria, enabling the development of a community with a high level of metabolic activity. A significant proportion of soil bacterial taxa exhibit a similar shift in life strategy: in the bulk soil (BS), they adopt an inactive or dormant state to survive the hostile environment, and in the rhizosphere (RZ), they resume metabolism and vegetative growth in response to root exudates during colonization. These findings underscore the importance of analyzing microbial community dynamics in harsher soil environments compared to typical croplands. Furthermore, correlating total and active taxa with spatial distribution data and physiological traits may improve our understanding of microbial seed bank dynamics.

## Acknowledgements

The authors thank Andrés Aravena for his valuable advice on data processing. PA thanks Scholarship INTA 2024T2DID for their support.

## Author contributions

D.E.P.: Conceptualization, Formal analysis, Methodology, Writing – original draft. C.S.: Conceptualization, Writing – original draft. A.G.: Investigation, Methodology. P.A.: Investigation. C.A.-C.: Investigation. C.A.-N.: Formal analysis. M.G.: Conceptualization, Funding acquisition, Methodology, Supervision, Project administration, Writing – original draft. V.C.: Conceptualization, Funding acquisition, Methodology, Project administration, Writing – original draft.

## Data availability

All sequencing data generated in this study have been submitted to the Sequence Read Archive (SRA) of the National Center for Biotechnology Information (NCBI) under BioProject accession number PRJNA1219082.

