## Supplementary figures for "The rhizosphere of *Pappostipa frigida* as a hotspot of active bacterial communities in the Andean steppe of the Atacama Desert"

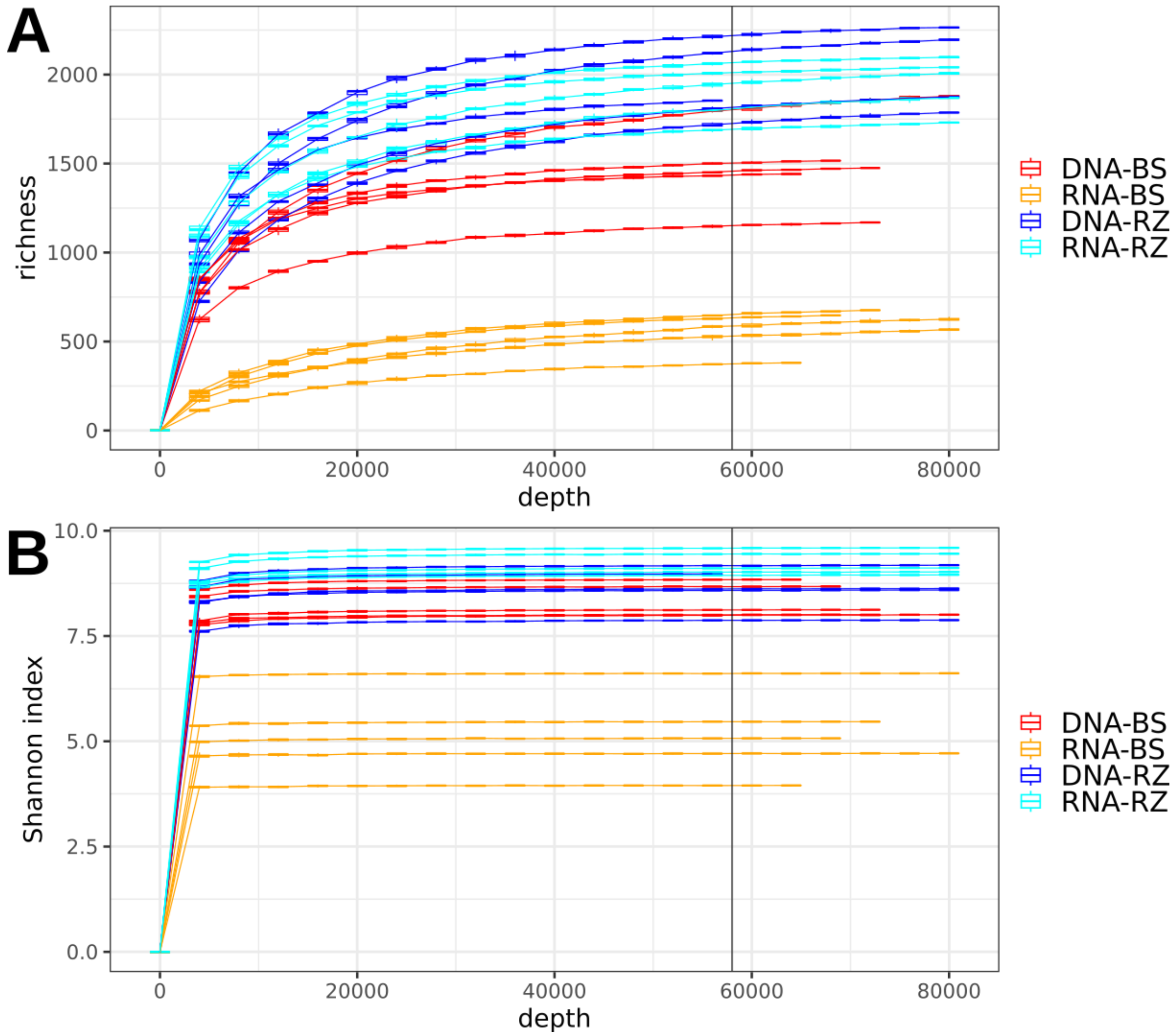


**Figure S1.** Rarefaction curves for (A) richness and (B) Shannon index.

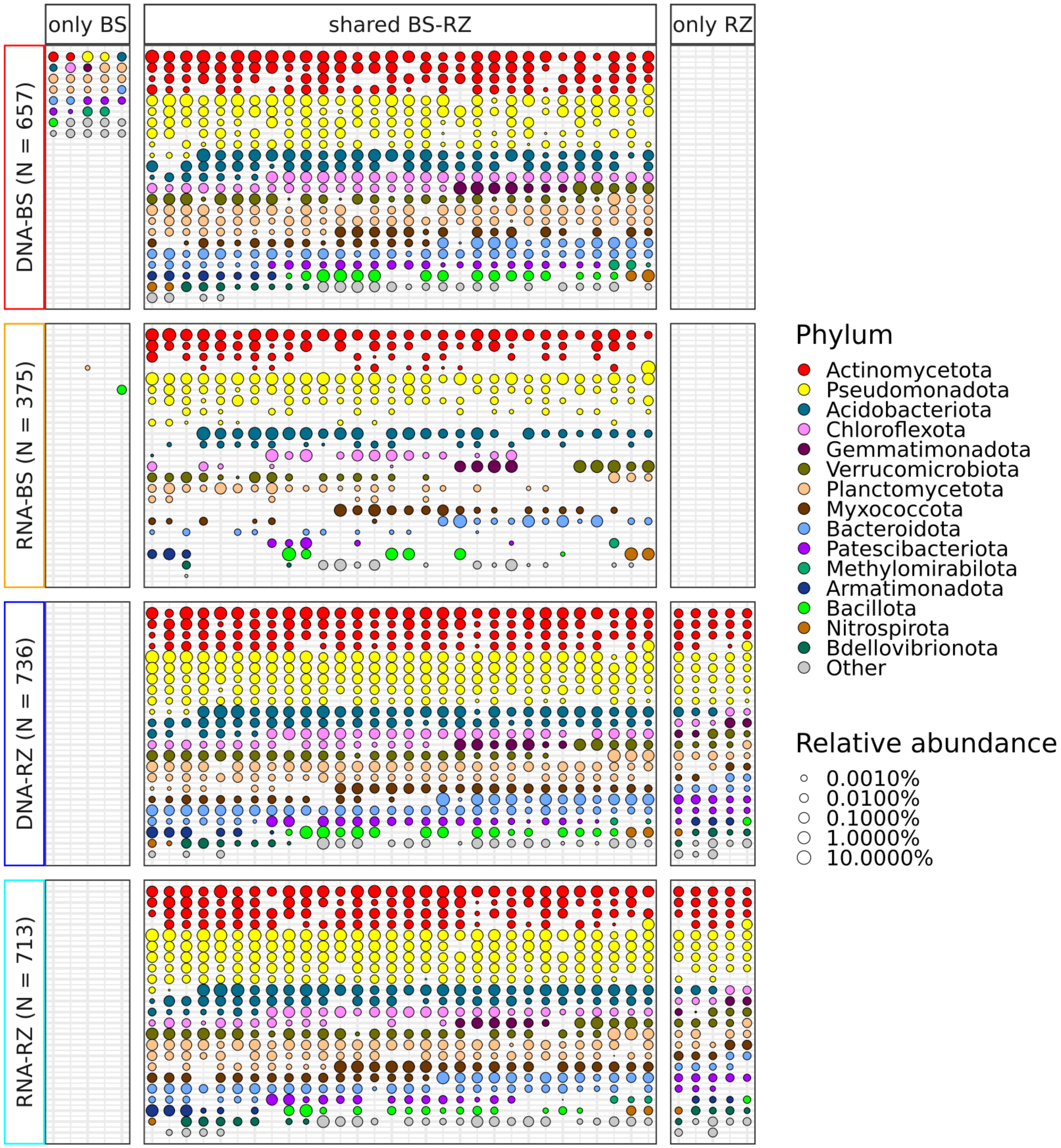


**Figure S2.** Composition of total (DNA) and active (RNA) bacterial genera in soil samples. The vertical panels categorize genera as unique to bulk soil (left, only BS), unique to the rhizosphere (right, only RZ), or shared between them (middle, shared BS-RZ). The horizontal panels display the different sample types (DNA-BS, RNA-BS,

DNA-RZ, RNA-RZ). For each sample type, all the detected genera are shown as dots, maintaining the same position in the panels of the other sample types, with colors indicating their phyla and sizes representing the log-transformed mean relative abundance in the respective sample type.

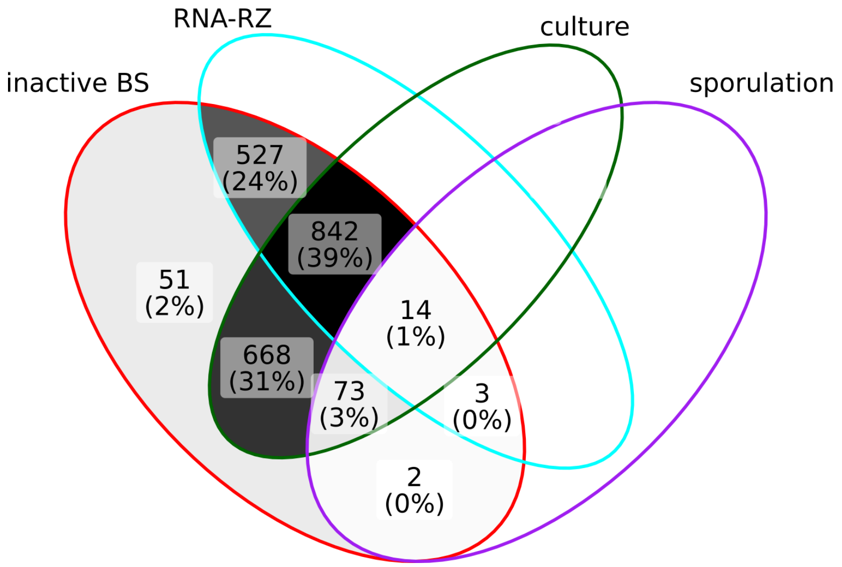


**Figure S3.** Inactive ASVs from BS shared with the RNA-RZ and culture (BS-SEM and BS-LB 10% NaCl) samples, as well as Bacillota with the capability for sporulation.
